# Identification of immunoreactive linear epitopes of *Borrelia miyamotoi*

**DOI:** 10.1101/741009

**Authors:** Rafal Tokarz, Teresa Tagliafierro, Adrian Caciula, Nischay Mishra, Riddhi Thakkar, Lokendra V Chauhan, Stephen Sameroff, Shannon Delaney, Gary P. Wormser, Adriana Marques, W. Ian Lipkin

**Affiliations:** Center for Infection and Immunity, Mailman School of Public Health, Columbia University, New York, NY; Department of Psychiatry, Columbia University, New York, NY; Division of Infectious Diseases, New York Medical College, Valhalla, NY; Laboratory of Clinical Immunology and Microbiology, National Institute of Allergy and Infectious Diseases, National Institutes of Health, Bethesda, Maryland.

## Abstract

*Borrelia miyamotoi* is an emerging tick-borne spirochete transmitted by *Ixodid* ticks. Current serologic assays for *B. miyamotoi* are impacted by genetic similarities to other *Borrelia* and limited understanding of optimal antigenic targets. In this study, we employed the TBD-Serochip, a peptide array platform, to identify new linear targets for serologic detection of *B. miyamotoi*. We examined a wide range of suspected *B. miyamotoi* antigens and identified 352 IgM and 91 IgG reactive peptides, with the majority mapping to variable membrane proteins. These included peptides within conserved fragments of variable membrane proteins that may have greater potential for differential diagnosis. We also identified reactive regions on FlaB, and demonstrate crossreactivity of *B. burgdorferi* C6 with a *B. miyamotoi* C6-like peptide. The panel of linear peptides identified in this study can be used to enhance serodiagnosis of *B. miyamotoi*.

## Introduction

*Borrelia miyamotoi* is a tick-borne relapsing fever spirochete found throughout temperate regions worldwide that is transmitted by hard ticks of the genus *Ixodes* (1-5). *B. miyamotoi* was discovered in 1995; however, the link to disease was first established in 2011 when it was implicated in an outbreak of tick-borne illness in Russia (1, 6). *B. miyamotoi* is primarily transmitted by *I. scapularis* and *I. pacificus* in North America, and *I. ricinus* and *I. persulcatus* in Europe and Asia (6-9). These tick species also transmit *Borreliae* that cause Lyme borreliosis (10). Lyme borreliosis is the most common vector-borne disease in the United States, where most infections are caused by *Borrelia burgdorferi* and transmitted by *Ixodes scapularis.* Despite sharing the same vectors, *B. miyamotoi* differs from *B. burgdorferi* in a number of ecological aspects, including the ability for transovarial transmission, quicker transmission during tick feeding, and lower infection rates in vector ticks. The prevalence of *B. miyamotoi* in nymphs is typically 1% to 5% versus 15% to 25% for *B. burgdorferi* (11-15). Nonetheless, *B. miyamotoi* and *B. burgdorferi* can occasionally infect the same tick, and concurrent human infections have been reported (5, 15, 16).

Symptomatic infections with *B. miyamotoi* (*Borrelia miyamotoi* disease; BMD) usually present with fever and other non-specific symptoms including fatigue, headache, chills and nausea (17, 18). Bouts of relapsing fever may occur in untreated patients. In the United States, reports of BMD are rare with less than 200 cases identified between 2011 and 2017 (3, 19, 20). This is considerably less than would be expected based on the high incidence of other tick-borne diseases and suggests a substantial underreporting of *B. miyamotoi* infections (21). A portion of infections may be asymptomatic while symptomatic infections may not be identified because of a similar presentation to certain other tick-borne illnesses and the lack of optimal diagnostic tests (22-24). Patients with BMD may test positive on the C6 ELISA, a serologic assay used in the diagnosis of Lyme disease (20, 25, 26). Current methods of BMD diagnosis include PCR (in the acute stage), and a two-tiered antibody assay (ELISA and western blot) based on immunoreactivity to glycerophosphodiester phosphodiesterase (GlpQ), an enzyme present in *B. miyamotoi* but absent in Lyme *Borrelia* (17, 27). Because GlpQ homologs are present in other relapsing fever spirochetes and in other bacteria, its specificity for *B. miyamotoi* is limited (28-31). In addition, serologic assays based on reactivity to GlpQ are only reactive in 56% to 78% of sera from convalescent BMD patients and only 16% of sera from individuals with acute disease (3, 32). Thus, other targets are needed in order to develop more sensitive and specific serologic tests.

An alternative approach for BMD serologic diagnosis is the development of assays that target outer surface antigens. Although this approach has been applied for diagnosis of Lyme borreliosis, relatively little is known about the utility of outer surface antigens in diagnosis of BMD. The primary challenge in assay development is the rapid antigenic variation that is a characteristic of relapsing fever spirochetes (33-36). Like all relapsing fever *Borreliae*, the genome of *B. miyamotoi* contains linear plasmids that encode multiple alleles of variable major proteins (Vmps) (36-38). Vmps consist of two types of lipoproteins that are dissimilar in sequence and length: the variable small proteins (Vsps) and variable large proteins (Vlps)(39). At any given time, only one Vmp is expressed by the spirochete when a single allele is copied into the Vmp expression site. Relapsing fever *Borreliae* alternate expression of these Vmps in order to evade the host immune response (31, 32, 38). The alleles are genetically heterogenous; thus, it is unlikely that host IgM raised against one Vmp will neutralize spirochetes expressing another Vmp. Despite high genetic diversity, Vmps share conserved regions that may also be a target of neutralizing antibodies. Identifying these regions could be useful for diagnosis, particularly early in disease (37, 39). In this study, we employed the Tick-borne disease Serochip (TBD-Serochip), a novel serologic peptide array platform, to examine the antibody response in human patients with BMD to a wide array of *B. miyamotoi* antigens. Our work identified linear epitopes that could be applied in the development of improved diagnostic tests (40).

## Materials and Methods

### TBD-Serochip

The TBD-Serochip is a peptide array that consists of approximately 170,000 12-mer linear peptides designed from the protein sequence of the primary antigens of *Anaplasma phagocytophilum*, *B. burgdorferi*, *B. miyamotoi*, *Babesia microti*, *Rickettsia rickettsii*, *Ehrlichia chaffeensis*, Powassan virus and Heartland virus (40). The 12-mer peptides tile the sequence of each antigen with an 11 amino acid (aa) overlap to the preceding 12-mer peptide in a sliding window pattern. Antigens selected for inclusion on the TBD-Serochip were all previously reported to elicit an antibody response in humans. For *B. miyamotoi*, the TBD-Serochip includes 12-mer peptides designed from a wide array of Vmp and non-Vmp antigens (**Table 1**). For each selected antigen, we downloaded every available homologous protein sequence from the NCBI protein database. Sequences were aligned and used to design 12-mer peptides, with redundant peptides excluded prior to synthesis. This approach resulted in the TBD-Serochip incorporating peptides to all genetic variants of every included antigen.

**Table 1.**
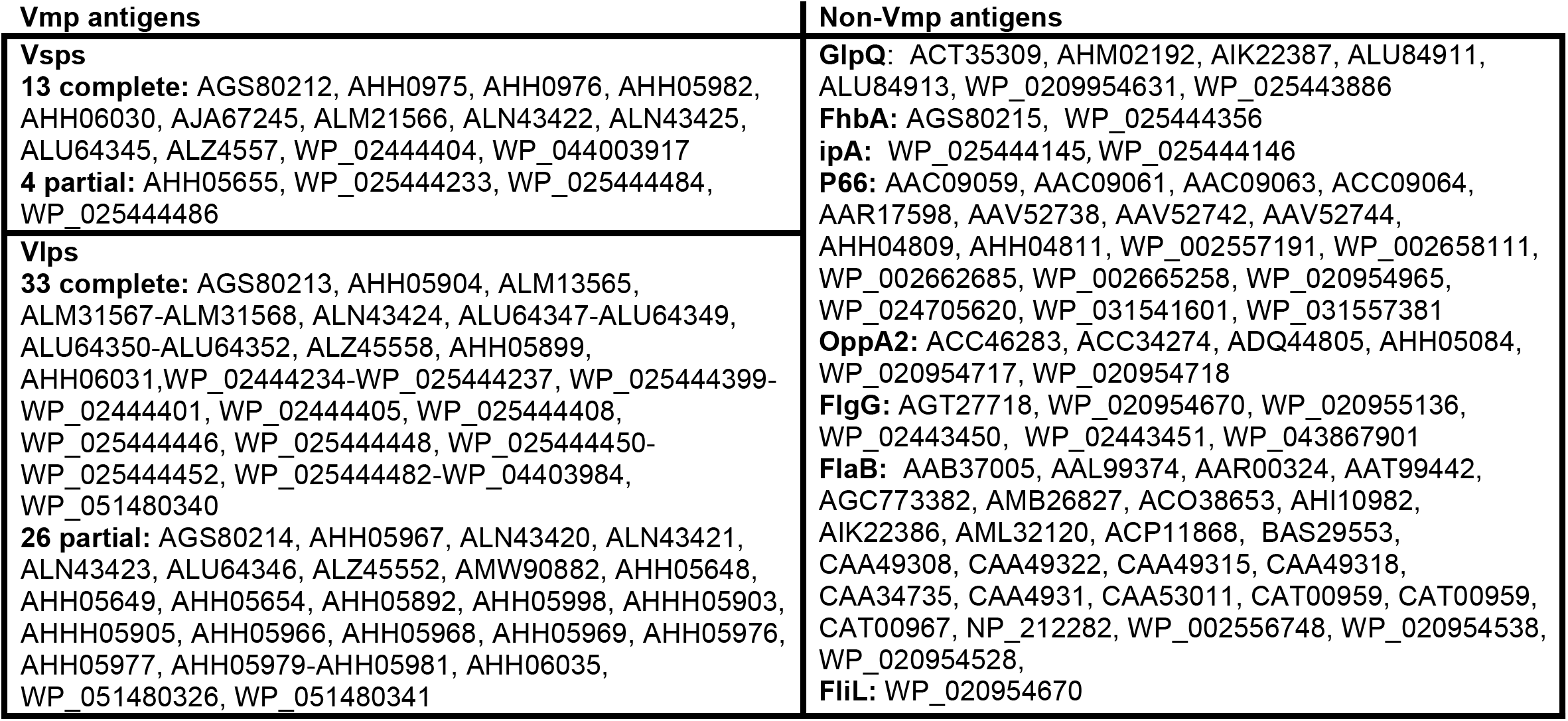
List of *B. miyamotoi* sequences used to design 12-mer peptides for the TBD-Serochip.

### Sera

We tested seven convalescent-phase sera from patients diagnosed with BMD. Five sera originated from New York Medical College, and 2 from Tufts University. Diagnosis of BMD was made during the acute disease phase, six by PCR, and one by a GlpQ-based serologic assay (**Table 2**). We contrasted these data with TBD-Serochip data obtained from sera from patients with Lyme disease (N=100), and healthy individuals with no history of Lyme disease (N=16). All sera were tested at a 1:50 dilution. We used de-identified human sera from our previous work with the TBD-Serochip, and additional samples acquired form the National Institutes of Health (40). These samples were obtained under clinical protocols (ClinicalTrials.gov Identifier <u>NCT00028080</u> and NCT00001539) approved by the institutional review board of the National Institute of Allergy and Infectious Diseases, and all participants signed informed consent. Sample types ranged from early to late Lyme disease (confirmed by the two-tiered testing algorithm), All methods used for sample collection were performed in accordance with the proper guidelines and regulations.

**Table 2.** BMD samples tested on the TBD-Serochip.

| Samples | BMD Diagnosis (acute stage) |
| --- | --- |
| BMD-1 | PCR |
| BMD-2 | PCR |
| BMD-3* | PCR |
| BMD-4 | PCR |
| BMD-5* | PCR |
| BMD-6 | PCR and GlpQ ELISA (IgM & IgG) |
| BMD-7 | GlpQ ELISA (IgM & IgG) |

\*previous diagnosis of Lyme disease

### Epitope identification

Following antibody incubations, the arrays were scanned on NimbleGen MS 200 Microarray Scanner (Roche) to generate raw fluorescent intensity data sets that were then subjected to quantile normalization. IgM and IgG reactivity were examined separately. Based on the data obtained from healthy controls, we computed a cut-off threshold for each epitope that was defined by calculating the mean plus 3 standard deviations of the signal intensity for the same epitope. We then filtered out any peptides with signal intensities below threshold in all experiments. To identify peptides that could be used to differentiate between BMD and healthy control sera, we used the non-parametric Wilcoxon rank sum test and identified peptides that yielded a statistically significant signal (q value ≤0.05 for both IgM and IgG, corrected for multiple testing using Benjamini-Hochberg Procedure). This identified the optimal sets of IgM and IgG 12-mer peptides that were reactive in sera with BMD, but not in healthy individuals. A portion of these 12-mer peptides were assembled into longer contigs that were then ranked by the average signal intensity.

### Luciferase ImmunoPrecipitation System assay (LIPS)

We generated a *Renilla* luciferase-GlpQ construct for detection of anti-GlpQ IgG antibody. The complete GlpQ coding sequence was amplified by PCR from a *B. miyamotoi-*positive *I. scapularis* using primers 5’-GGATCCATGAAATTAAAATTACTAATGC-3’ and 5’-GAGCTCTTATTTTTTTATGAAGTTCATT-3’. The PCR product was cloned into pREN2 vector and the fusion proteins were generated in Cos-1 cells. Light units were measured in a Berthold LB 960 Centro microplate luminometer (Berthold Technologies) using a Renilla Luciferase Assay System kit (Promega) and MikroWin 2010 software.

## Results

Of the 23,946 *B. miyamotoi* 12-mer peptides present on the TBD-Serochip, we identified 1491 peptides that were significantly reactive with IgM antibodies present in the seven BMD sera tested. Through assembly of overlapping reactive 12-mer peptides, we identified 352 reactive epitopes made up of 2 or more consecutive reactive peptides. For IgG, we identified 429 peptides that were clustered into 91 putative epitopes. All IgM and IgG epitopes were then mapped to specific regions within *B. miyamotoi* proteins. The majority of the epitopes mapped to Vmps (**Table 3**).

**Table 3.**
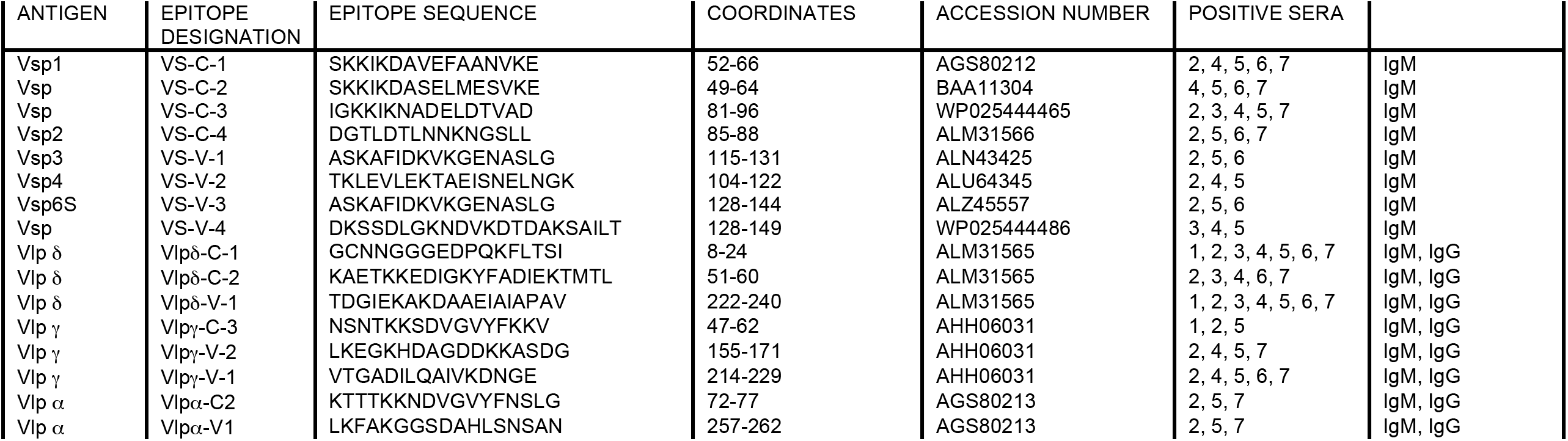
Major immunoreactive linear regions of *B. miyamotoi* Vmps identified with the TBD-Serochip.

In addition to *B. miyamotoi* peptides, we examined the reactivity to peptides from other tick-borne agents. In 2 samples, we detected IgG reactivity to specific epitopes of *B. burgdorferi* that were identified in our previous work (40). Subsequent examination of clinical history of these 2 individuals revealed that both had previously been diagnosed with Lyme disease.

### Reactivity to Vsp peptides

The TBD-Serochip contains 12-mer peptides designed from the amino acid (aa) sequence of 13 full length and 4 partial Vsp paralogs (**Table 1**). *B. miyamotoi* Vsps are approximately 220 aa in length and consist of a conserved N-terminal fragment of approximately 85 aa, followed by highly variable regions (**Figure 1A**). We identified IgM-reactive epitopes within both the conserved and variable fragments within every Vsp homolog. Although previous studies that were focused on Vsp1 suggested that it may be a major immunodominant antigen of *B. miyamotoi*, our data revealed Vsp4 to be more immunoreactive (**Figure 1B** and **1C**). Among the immunodominant regions of Vsps included a region within the N-terminal fragment that was reactive with IgM in 5 out of 7 sera (sera 2 and 4 - 7). This 16 aa peptide, designated VS-C-1, spans aa 52 to 66 of Vsp1 (mapped to accession number AGS80212) and is conserved in the majority of sequenced *B. miyamotoi* Vsp paralogs (**Figure 2**).

**Figure 1.**
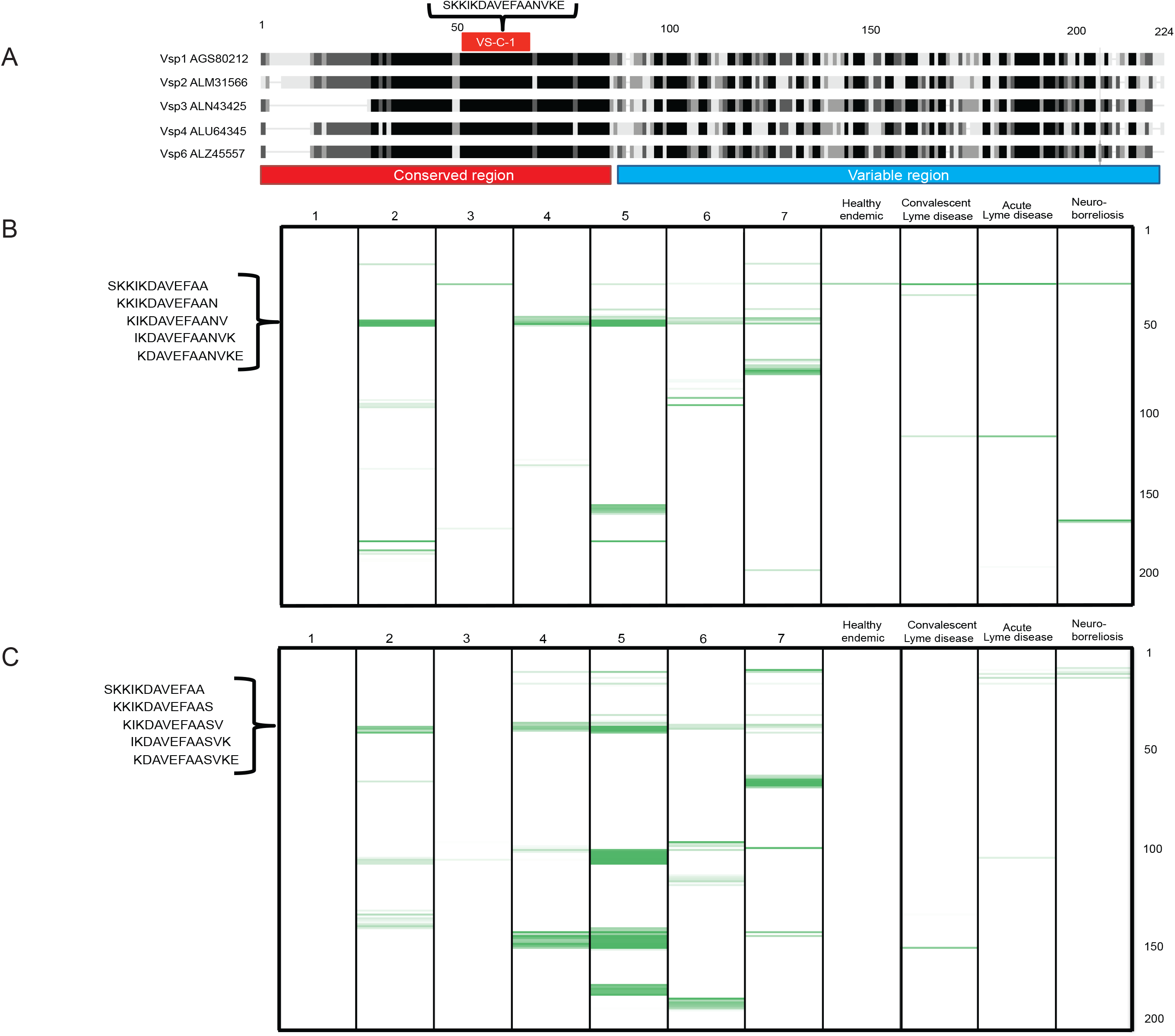
Identification of reactive epitopes within the conserved region of Vsps. Panel A displays the alignment of five *B. miyamotoi* Vsp homologs. Regions of homology are shown in black. The numbers on top of the alignment denote the relative aa position of each protein relative to Vsp1. The location of the consensus reactive epitope VS-C-1 and its aa sequence are indicated. Panels B and C show IgM reactivity plots of Vsp1 (accession number AGS80212) and Vsp4 (accession number ALU64345), respectively. Numbers 1 to 7 represent the BMD samples. Reactivity to control sera (from a healthy individual and various stages of Lyme disease) are shown on the right. The Y axis represents the location of 12-mer peptides positioned along the contiguous protein sequence of Vsp1 and Vsp4. Immunoreactivity with the 12-mer peptides is indicated in green, with darker color corresponding to increasing reactivity. The location of VS-C-1 (samples 2, 4, 5, 6 and 7) and its corresponding 12-mer peptide sequences are indicated in the bracket on the left in panels B and C.

**Figure 2.**
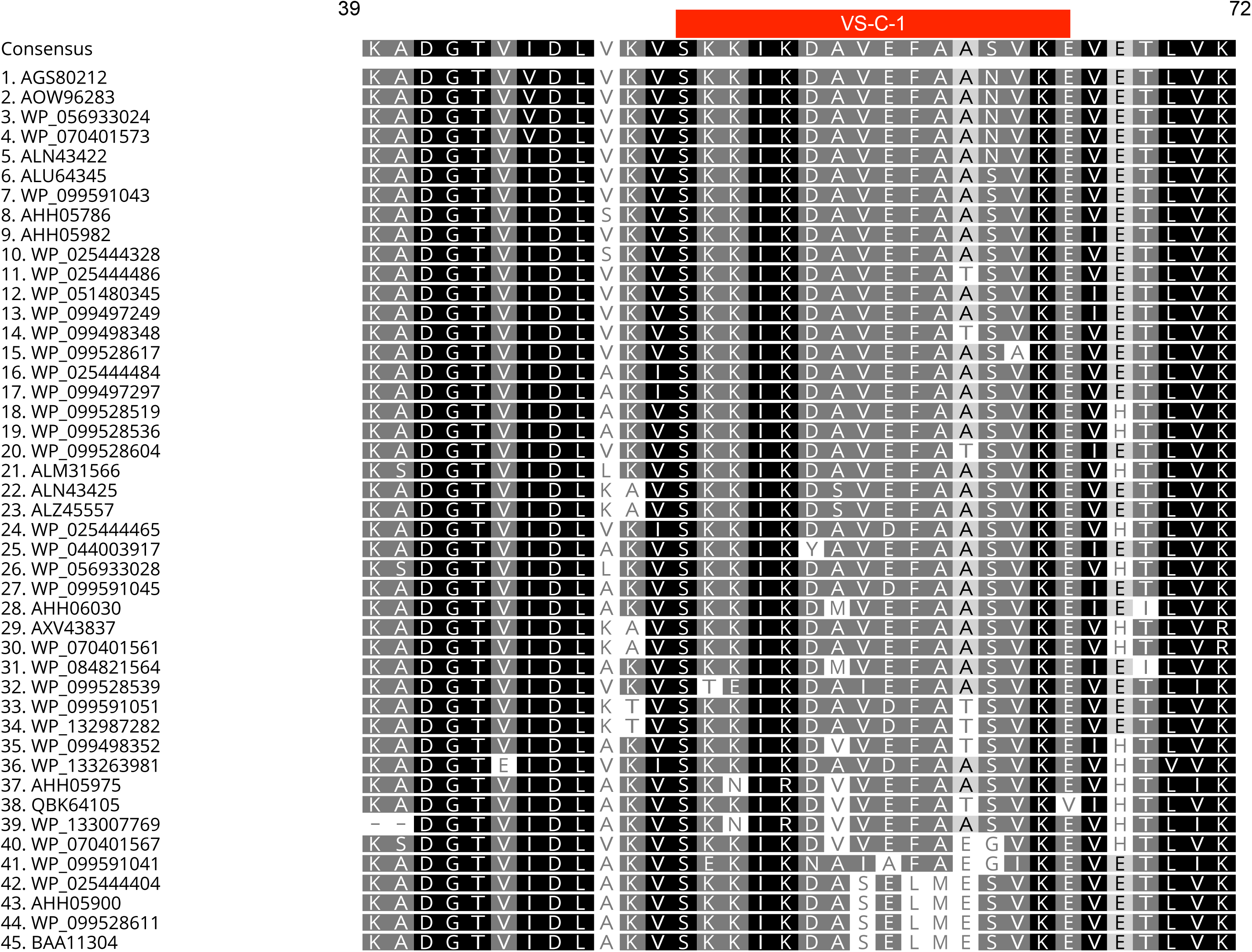
Conservation of the VS-C-1 peptide in *B. miyamotoi*. Shown is the aa alignment of all Vsp sequences deposited in Genbank as of June 28, 2019. Accession numbers are indicated on the left. Location of the VS-C-1 peptide is shown in red. The numbers on top denote the aa position within Vsp1.

One Vsp paralog (accession number BAA11304) contains substantial aa variation within the central portion of the VS-C-1 sequence. The aa at positions 8-12 in this paralog deviates from AVEFA to SELME (**Figure 2**). Presumably, antibodies to VS-C-1 region would not react with peptides with this divergent amino acid sequence (designated VS-C-2). Nonetheless, peptides corresponding to VS-C-2 (mapped to accession number BAA11304) were reactive with 4 of the 7 VS-C-1-reactive samples (sera 4-7), but did not react with sera from patient 2 (**Figure 3**). The presence of antibodies to both peptide variants indicates that these 4 patients were exposed to Vsp paralogs with both VS-C-1 and VS-C-2 sequences.

**Figure 3.**
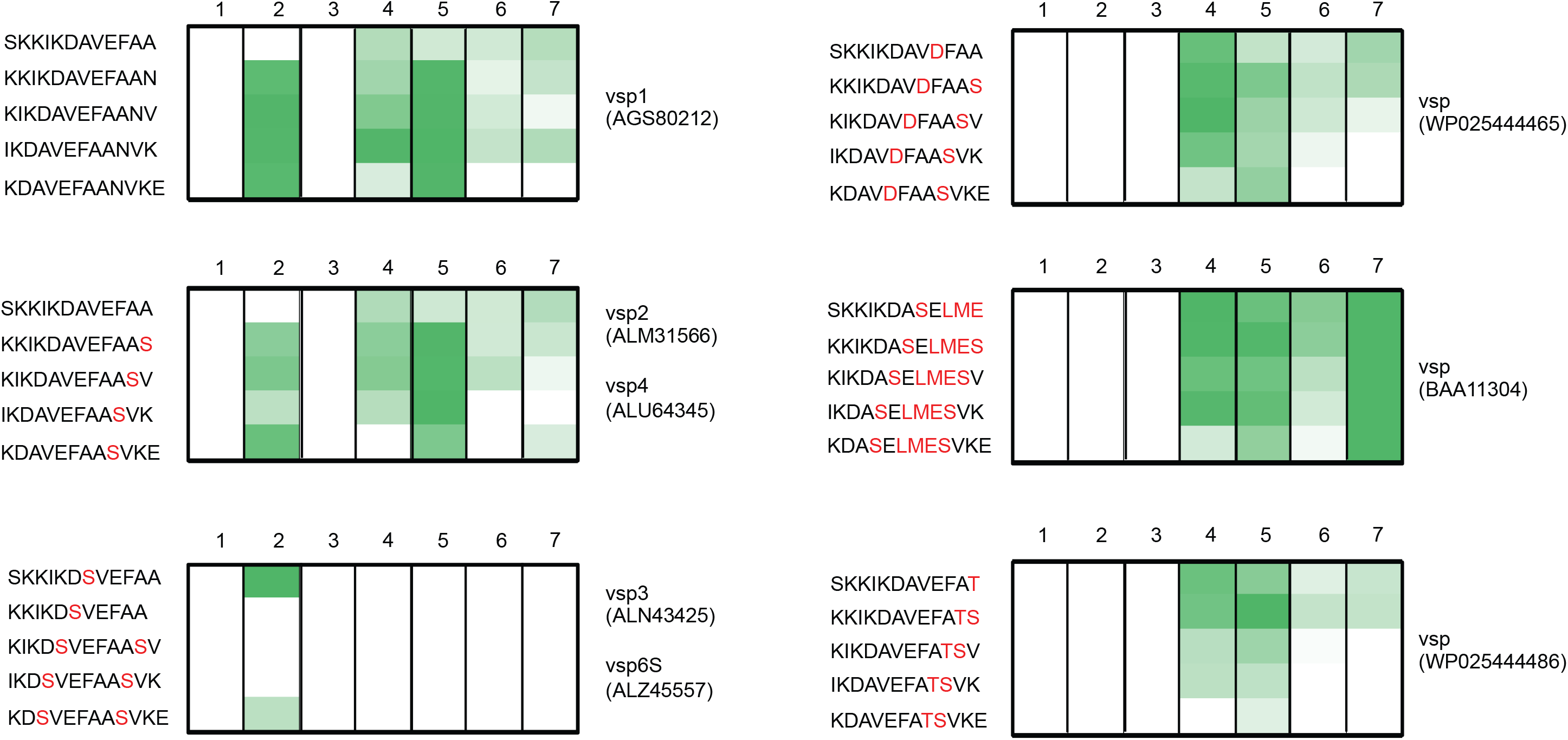
Vsp diversity and reactivity to the Vsp-C-1 epitope. Each panel represents the reactivity of the seven tested sera (labeled 1-7) to the peptides from Vsp homologs present on the TBD-Serochip. The variant Vsp 12-mer peptide sequences are shown on the left; accession numbers are indicated on the right. Amino acid differences to the consensus sequence (based on Vsp1) are displayed in red. Immunoreactivity to the 12-mer peptides is indicated in green with increasing signal intensity displayed from light to dark.

We identified other reactive regions on Vsps, including VS-C-3 located at the end of the conserved N-terminal portion that was highly reactive in sera from patient 7 (**Figure 1B**). We also identified epitopes that were mapped to the variable fragments of Vsps. However, these epitopes were less frequently reactive than VS-C-1 or VS-C-2 (**Table 3**). OspC, a highly immunogenic *B. burgdorferi* lipoprotein, is the closest homolog to Vsp. Both proteins have a high degree of homology within the N terminal region. We and others previously identified a reactive epitope within this region of OspC (39, 41, 42). The alignment of Vsp1 (AGS80212) with OspC (CAA59253) revealed that the VS-C-1 and OspC epitopes do not overlap and map to non-homologous portions of these lipoproteins (**Figure 4**).

**Figure 4.**
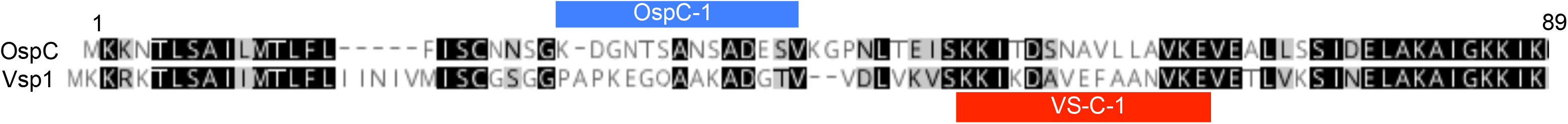
Locations of major immunogenic linear peptides in Vsp and OspC. The N terminal portions of Vsp1 and OspC were aligned to contrast the reactive portions of OspC (in blue) and Vsp (in red).

### Reactivity to Vlp peptides

Vlps of relapsing fever *Borrelia* are typically 330 aa to 350 aa in length and are classified into α, β, δ, and γ sub-families (38). The N-terminal 70 aa to 120 aa fragments are conserved within each subfamily, whereas the remaining portions display a substantially greater heterogeneity in aa sequence. *B. miyamotoi* encodes multiple α, δ, and γ alleles and only a single putative β-like allele. To account for this genomic diversity, the TBD-Serochip includes peptides for 33 full length and 26 partial α, δ, and γ homologs from multiple *B. miyamotoi* strains (**Table 1**).

All 7 tested sera reacted with a wide range of peptides from Vlps. Although IgM reactivity was predominant, several Vlp regions were also reactive with IgG. Overall, the largest number of Vlp-reactive peptides mapped to δ Vlps. **Figure 5** displays the location of all reactive peptides mapped to Vlp15/16 (accession number ALM31565), a Vlp homolog with the highest number of immunoreactive epitopes. A δ Vlp-specific region, located between aa 8-24 of Vlp15/16, reacted with all seven BMD sera (**Figure 5**). We designated this region Vlpδ-C-1 (**Table 4**). Six sera were IgM-positive to Vlpδ-C-1, including two (samples 1 and 3) that were also IgG-positive. One serum (sample 2) was reactive with only IgG. Six sera were reactive with another reactive region designated Vlpδ-C-2. Two sera were reactive with IgM and IgG, two with IgM, and two with IgG. Vlp-C-2 was mapped to aa 51-66 of Vlp15/16, a fragment that is also partially conserved in α, δ, and γ Vlps. γ Vlps peptides corresponding to this region (designated Vlpγ-C-2) were reactive with four sera (all IgG) (**Figure 6, Table 3**). α peptides (Vlpα-C2) were reactive with IgG from three sera (**Table 3**).

**Table 4.** Reactivity of the C6 peptides in *B. burgdorferi* and *B. miyamotoi*.

| BMD samples | Prior Lyme disease | Reactivity |  |  |  |
| --- | --- | --- | --- | --- | --- |
|  |  | TBD-Serochip |  | C6 ELISA | Cross reactivity on the C6 ELISA |
|  |  | B. miyamotoi C6-like | B. burgdorferi C6 |  |  |
| 1 | - | - | - | - | - |
| 2 | + | + | + | + | + |
| 3 | - | - | + | + | - |
| 4 | - | - | - | - | - |
| 5 | + | + | + | + | - |
| 6 | - | + | + | + | + |
| 7 | - | + | + | + | + |

+ indicates reactivity to the 12-mer peptides within the C6 or C6-like region

**Figure 5.**
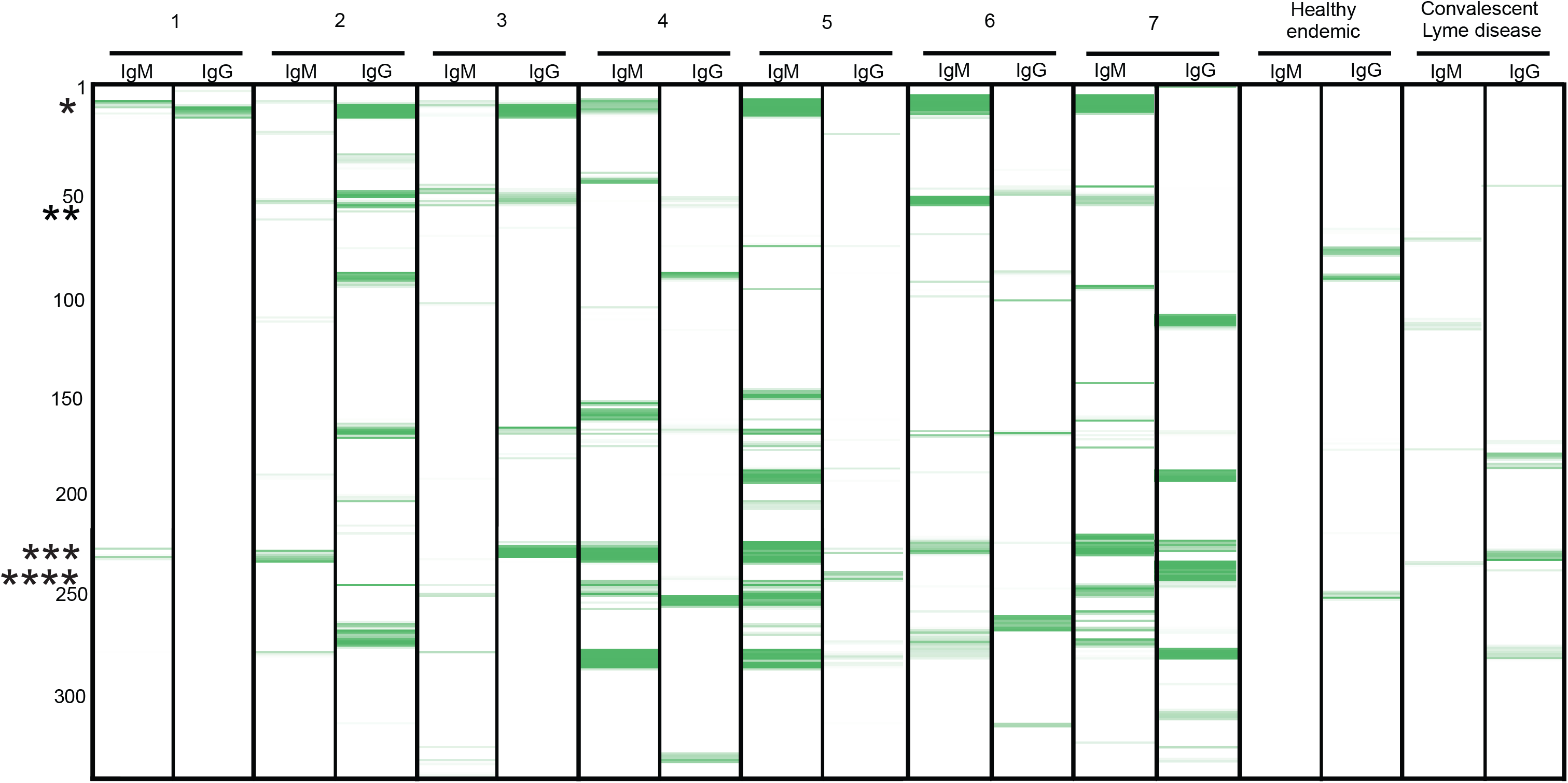
Identification of reactive peptides within Vlp δ. Shown is the IgM and IgG reactivity map displaying reactive 12-mer peptides (in green) of Vlp 15/16 (accession number ALM31565). Numbers 1 through 7 represent the 7 BMD sera. Reactivity to control sera (healthy individual from a Lyme endemic area and a patient with Lyme disease at convalescence) are shown on the right. The numbers on the Y-axis represent the aa location of the 12-mer peptides positioned along the contiguous protein sequence of Vlp 15/16. Regions with immunoreactive 12-mer peptides are indicated in green with increasing signal intensity displayed from light to dark. The asterisks indicate major reactive epitopes; * Vlpδ-C-1, ** Vlpδ-C-2, *** Vlpδ-V-1, **** C6-like.

**Figure 6.**
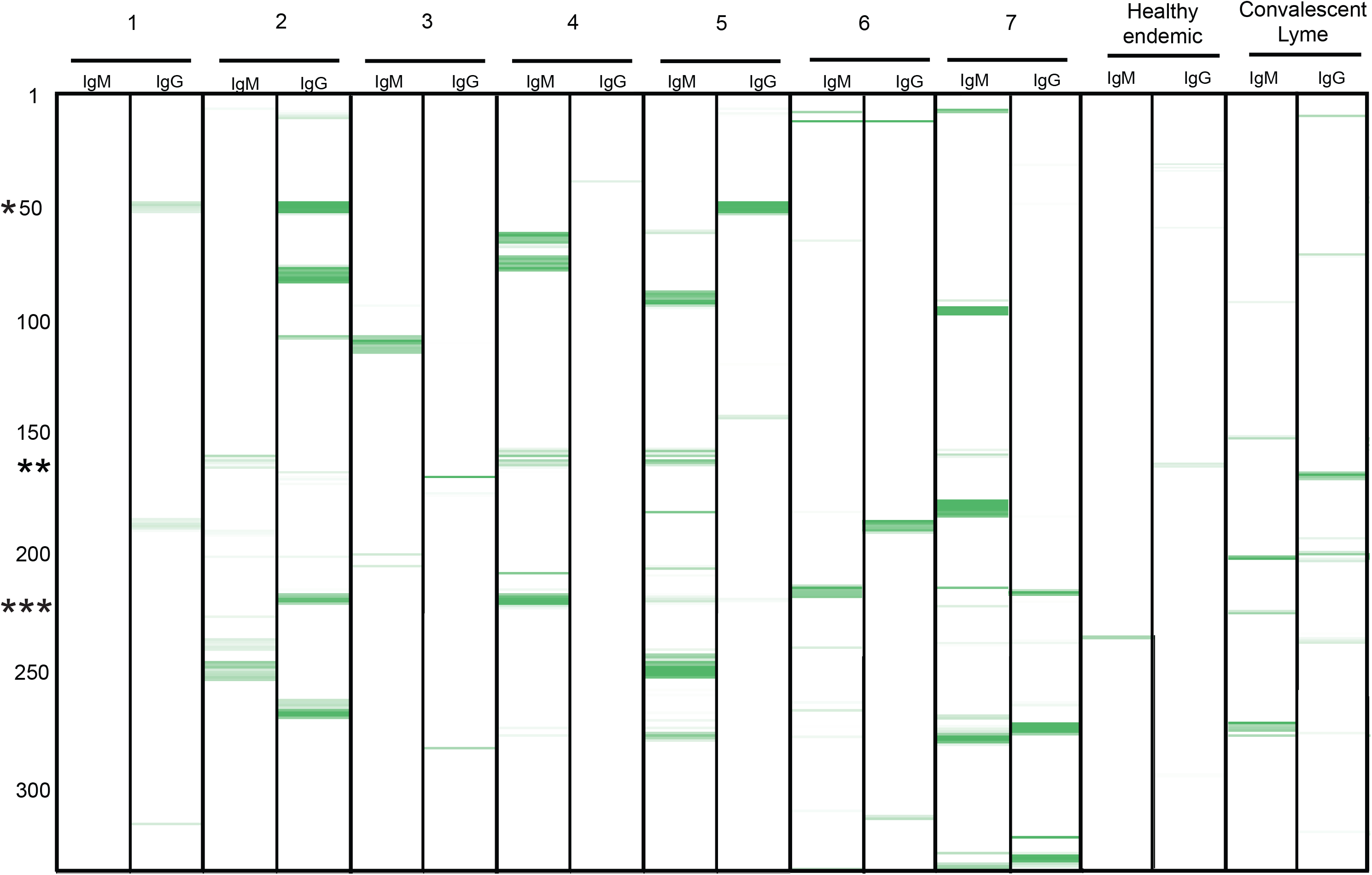
Identification of reactive peptides within Vlp γ. Shown is the IgM and IgG reactivity map displaying reactive 12-mer peptides (in green) of Vlp5 (accession number AHH06031). Numbers 1 through 7 represent the 7 BMD sera. Reactivity to control sera (healthy individual from a Lyme endemic area and a patient with Lyme disease at convalescence) are shown on the right. The numbers on the Y-axis represent the aa location of the 12-mer peptides positioned along the contiguous protein sequence of Vlp 5. Regions with immunoreactive 12-mer peptides are indicated in green with increasing signal intensity displayed from light to dark. The asterisks indicate major reactive epitopes; * Vlpγ-C-3, ** Vlpγ-V-2, *** Vlpγ-V-1

Only one region within the variable fragment of δ Vlps was reactive with all seven sera. This region, mapped to aa 222-240 of Vlp 15/16, was designated VlpδV-1. In three sera, the reactive fragment extended approximately 25 aa downstream, and overlapped with a conserved region that is homologous to the C6 peptide of *B. burgdorferi* (**Figure 7A**). VlpδV-1 corresponds to a poorly conserved fragment in Vlps, even within paralogs of the same subfamily (**Figure 7A**). Nonetheless, all seven sera were reactive to a wide range of non-homologous peptides from different δ Vlp homologs that mapped to this region (**Figure 7B**). Although the corresponding peptides from Vlp α and Vlp γ have limited homology to VlpδV-1 peptides from Vlp δ, 4 sera also had reactivity to Vlp γ peptides and 3 sera reacted with Vlp α peptides from within this region. Our results indicate that the region corresponding to Vlp-V-1 in Vlps represents a major immunogenic region within *B. miyamotoi* Vlps, irrespective of sequence similarity.

**Figure 7.**
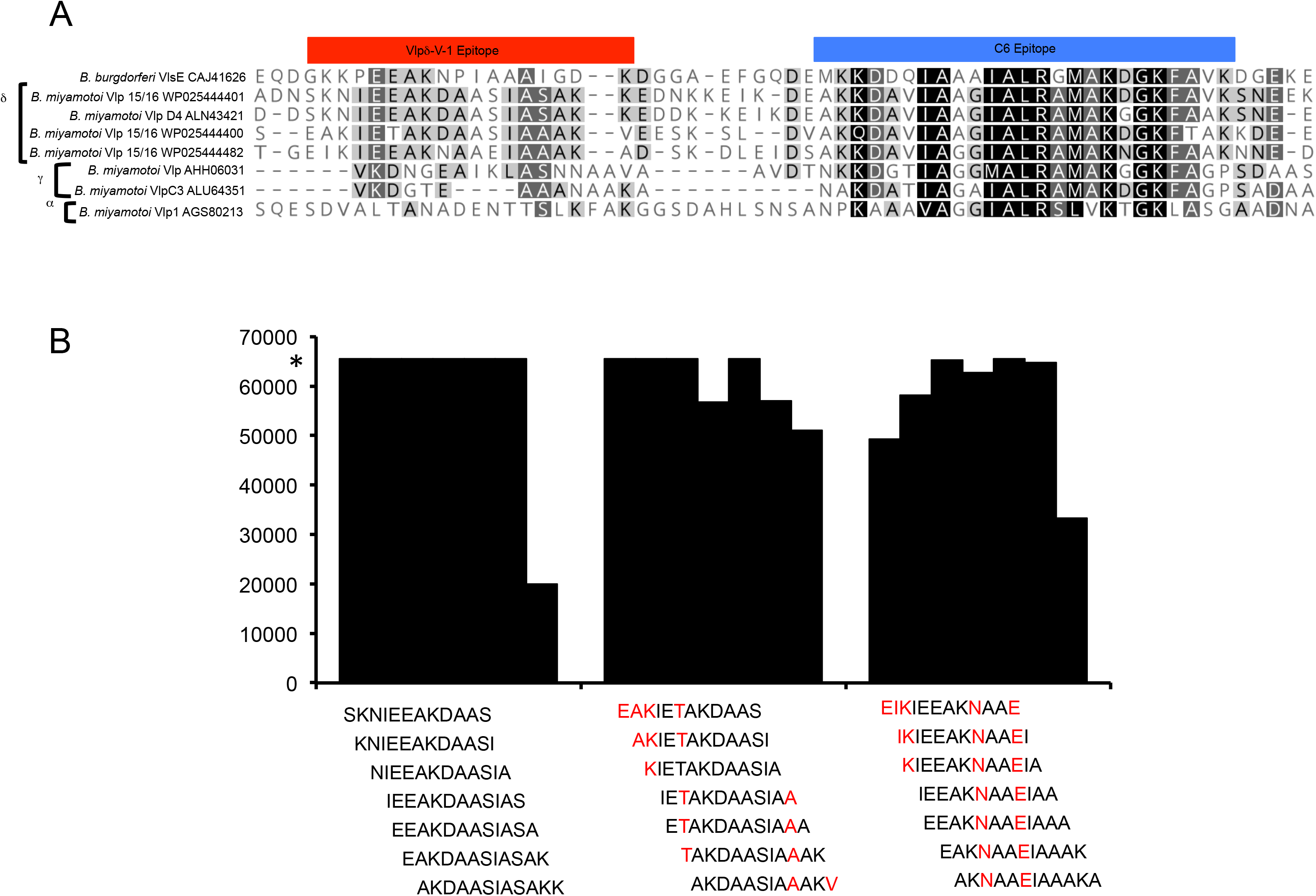
Sequence heterogeneity does not impact the reactivity to Vlpδ-V-1. Panel A - alignment of *B. miyamotoi* Vlp homologs with *B. burgdorferi* VlsE. The position of Vlpδ-V-1 reactive region is indicated in red, and the *B. burgdorferi* C6 epitope in blue. Panel B - Signal intensity of the 12-mer peptides representing Vlpδ-V-1 sequence variants. * - maximum intensity.

### Crossreactivity of the C6 peptides

VlsE, the closest homolog to Vlp, is a major immunodominant *B. burgdorferi* lipoprotein and includes a 26 aa C6 epitope that is employed in a peptide ELISA for Lyme disease diagnosis. Recent studies have reported cross-reactivity in the Lyme disease C6 ELISA with sera from BMD patients (20, 26). The similarity between a C6-like region in *B. miyamotoi* and the C6 was cited as a potential cause. Comparison of *B. burgdorferi* C6 and corresponding homologous Vlp sequences indicate that some δ Vlp15/16 homologs share 20 out of 26 aa residues with the C6 epitope (**Figure 7A**). The homology is most pronounced at the C terminal portion, with 14 out of 15 identical residues in both peptides. To more clearly delineate antibody responses to these two fragments, we compared the TBD-Serochip data to results obtained with a commercial C6 ELISA. Of the seven BMD sera tested, four samples (2, 5, 6, and 7) were positive on the TBD-Serochip for reactivity to the *B. miyamotoi* C6-like peptides (**Figure 8**). The same 4 samples, along with sample 3, had reactivity to the *B. burgdorferi* C6 peptides and all were also positive on the C6 ELISA. Sera 1 and 4 were negative with both assays. Serum 3 was positive for *B. burgdorferi* with the C6 ELISA but did not react with *B. miyamotoi* peptides on the TBD-Serochip. This discrepancy was likely due to past exposure to *B. burgdorferi*. Samples 3 and 5 both contained IgG antibodies to the C6 peptides from a prior infection with *B. burgdorferi.* These antibodies were specific for *B. burgdorferi* peptides and did not crossreact with the corresponding peptides of *B. miyamotoi* **(Figure 8)**. Serum 3 did not contain antibodies to the *B. miyamotoi* C6-like region; thus the positive result on the *B. burgdorferi* ELISA and TBD-Serochip result was likely exclusively triggered by anti-*B. burgdorferi* antibodies. Serum 5 had IgG antibodies to *B. burgdorferi*, and IgM antibodies to *B. miyamotoi* C6-like peptides. Sera 2, 6 and 7 had IgM or IgG antibodies to the *B. miyamotoi* C6-like peptides; they also reacted with the highly conserved C-terminal portion of the *B. burgdorferi* C6 and were the likely cause of the positive result on the C6 ELISA.

**Figure 8.**
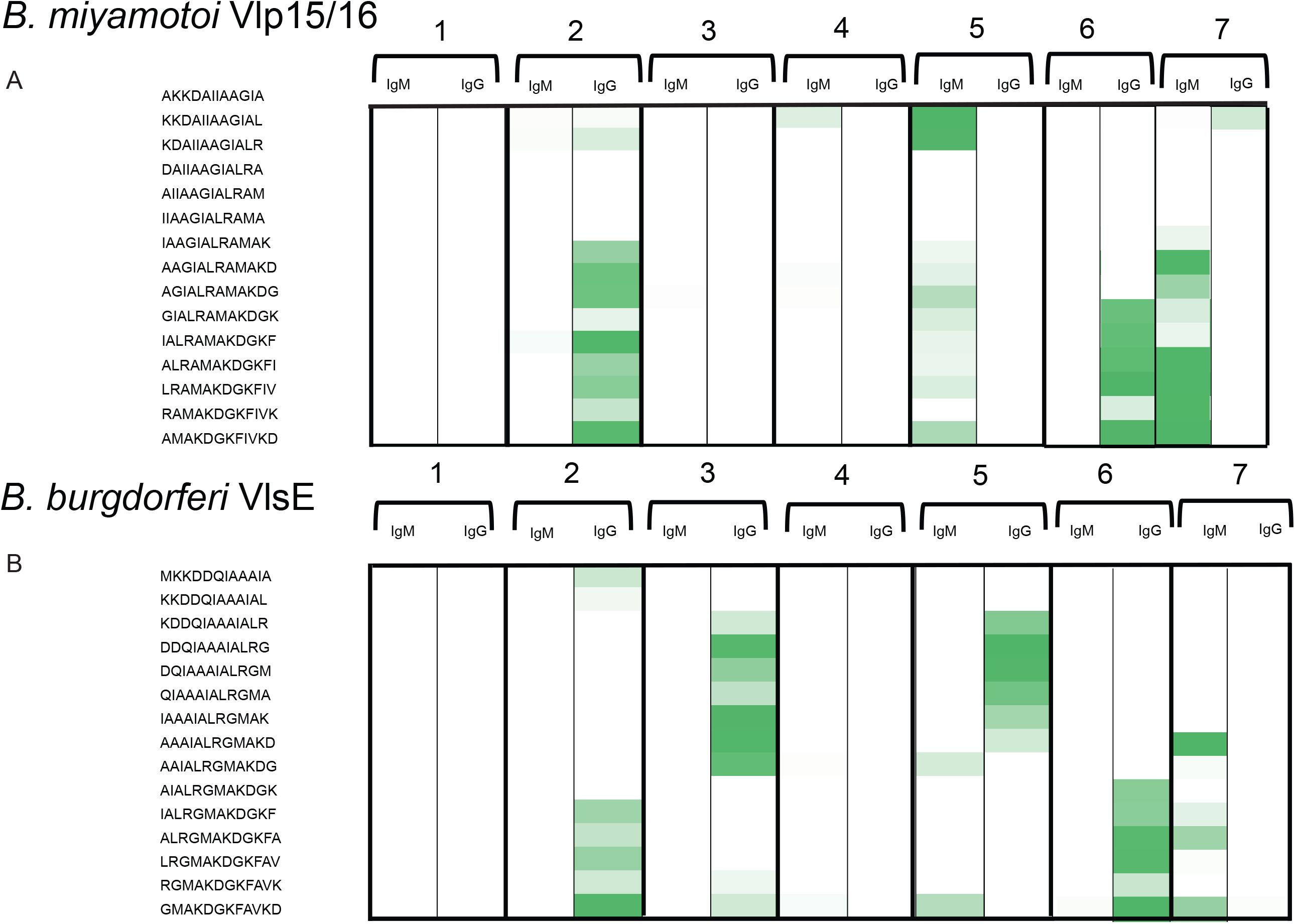
Immunoreactivity comparison of the *B. burgdorferi* C6 and the C6-like peptide of *B. miyamotoi*. Panel A displays the reactivity maps of the 12-mer peptides that constitute the 26 aa C6-like region of *B. miyamotoi*. Panel A shows the corresponding C6 12-mer peptides from *B. burgdorferi*. The individual 12-mer peptides are indicated on the Y axis. Regions with immunoreactive 12-mer peptides are displayed in green with increasing signal intensity displayed from light to dark.

In summary, only 4 of the 7 patients with a history of *B. miyamotoi* infection had antibodies to the *B. miyamotoi* C6-like peptide. This is in contrast to the C6 which is one of the most frequently reactive linear peptides in *B. burgdorferi.* However, when they are present, the antibodies to the *B. miyamotoi* C6-like peptide will likely crossreact on the C6 ELISA.

### FlaB

FlaB was the most reactive non-Vmp antigen. IgM antibodies to FlaB were detected in six out of seven BMD sera tested. FlaB comprises the major component of the spirochete flagellum and is among the most immunogenic of all *Borrelia* antigens (43, 44). However, its utility in diagnosis is compromised by its high cross-reactivity (45). FlaB bands can be recorded on both IgM and IgG Lyme disease western blots even in specimens from healthy individuals (46). FlaB is also highly conserved amongst *Borrelia,* with *B. miyamotoi* and *B. burgdorferi* FlaB sharing 90% aa identity. This further limits its utility for differential diagnosis. In tests of BMD or Lyme disease sera, we detected reactivity to a wide range of corresponding FlaB 12-mer peptides from both *B. miyamotoi* and *B. burgdorferi* (**Figure 9 A and B**). The same was true when we tested sera from patients with Lyme disease. We mapped the primary reactive portion of FlaB to a 45 aa fragment located between aa 192 and 236 of *B. miyamotoi* (accession number AHH05270) and its corresponding region in *B. burgdorferi* located between aa 190 and 236 (accession number AAC66541) (**Figure 9 C**). We found this region to be among the most frequently reactive peptide fragments when testing sera from Lyme disease or BMD. Despite this high reactivity, the majority of 12-mer peptides within these fragments were occasionally cross-reactive with IgM antibodies present in sera from healthy individuals. Nonetheless, we identified a 14 aa fragment within this region that was reactive only with BMD and Lyme disease sera but not with control sera. This peptide, AQEGAQQEGVQAVP, was located within aa 210 and 223 of *B. miyamotoi* FlaB. The corresponding 13 aa peptide in *B. burgdorferi* VQEGVQQEGAQQP was located within aa 211 and 223. Although these peptides cannot be used to discriminate between *B. miyamotoi* and *B. burgdorferi*, they may have utility for diagnosis of *Borrelia* infections.

**Figure 9.**
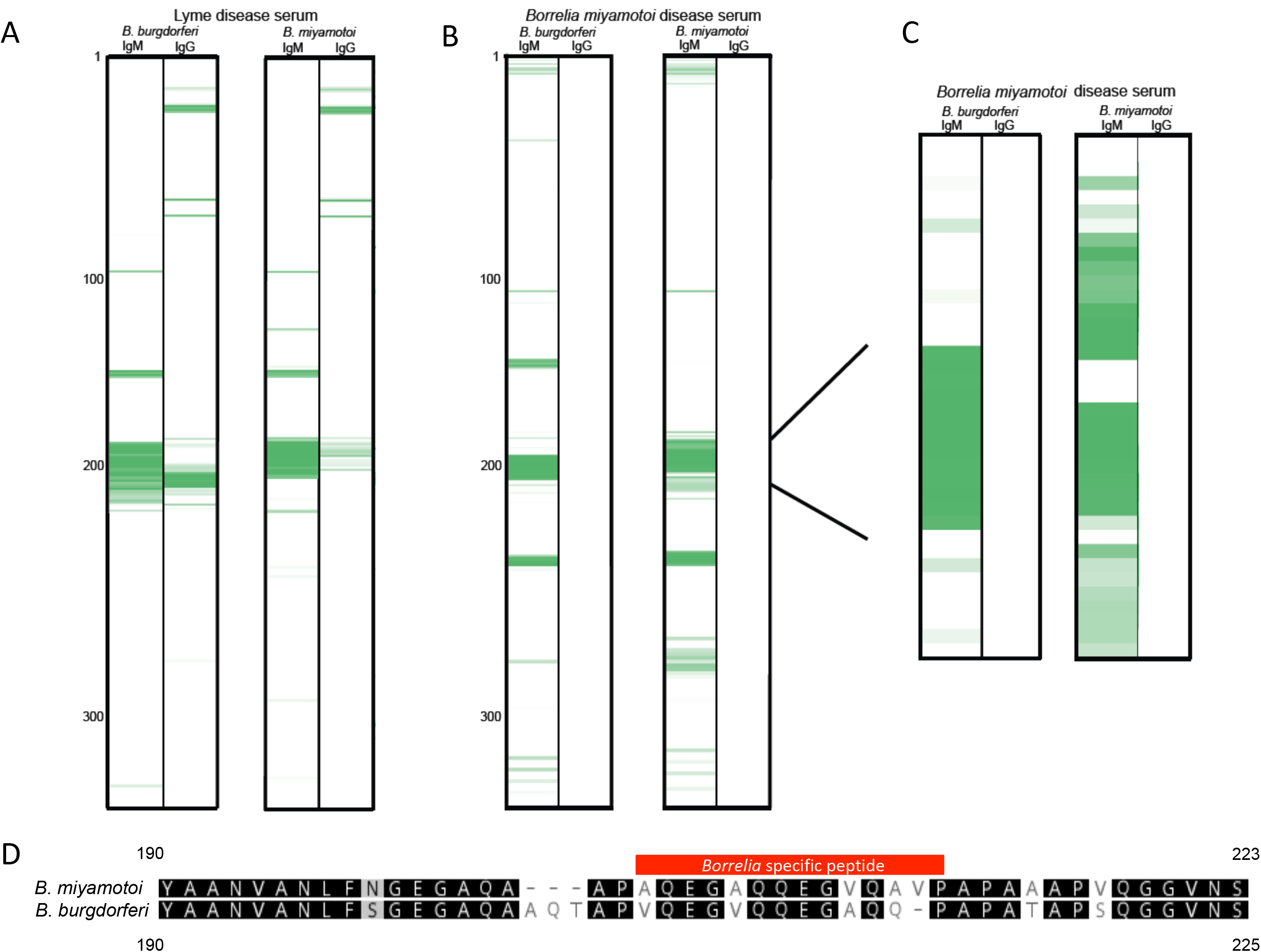
Reactivity to FlaB. Reactivity plots from a convalescent-phase Lyme disease serum (panel A) and BMD serum (Panel B). The reactive regions (in green) are indicated on the contiguous protein sequence of *B. burgdorferi* (accession number AAB36994-left) and *B. miyamotoi* (accession number AAT99442-right). Panel C shows a close up of the primary reactive region, located between 190-220 on both antigens. Panel D displays an alignment of the primary reactive region, with the *Borrelia* specific epitope indicated in red.

### GlpQ

We did not identify a specific epitope for GlpQ. To determine whether this was due to the absence of anti-GlpQ antibodies, we established a LIPS assay using a full length GlpQ as a target antigen. We tested 5 samples and IgG antibodies were present in 3 sera. Our combined TBD-Serochip and LIPS data suggests that these 3 samples contained anti-GlpQ antibodies to conformational but not linear epitopes. We propose that detection of antibodies to conformational epitopes likely constitutes the primary means of GlpQ serologic detection.

We did not identify unique epitopes on the remaining non-Vmp antigens. Although reactive peptides were detected, they were not consistently reactive among the samples tested.

### Additional specimens

We analyzed three additional sera from patients who had received a BMD diagnosis based on a positive index on a GlpQ ELISA from a commercial laboratory. Upon testing these specimens on the TBD-Serochip, we did not observe reactivity to any *B. miyamotoi* antigens. When we examined these by our LIPS assay, one sample had a very low positive reading and the remaining two were negative.

## Discussion

Lack of standardized assays, coupled with limited understanding of optimal target antigens contribute to the challenge of serologic diagnosis of TBD (47-49). The identification of superior targets, particularly of immunodominant specific epitopes has the potential to improve serodiagnosis. Among the primary challenges of differential serologic diagnosis of BMD is the insufficient sensitivity of GlpQ and the antigenic similarities between *B. miyamotoi* and Lyme borreliosis *Borrelia*. Through accurate mapping of specific linear immunoreactive peptides, the TBD-Serochip provides an unparalleled opportunity for identification of agent-specific linear epitopes that could facilitate differential diagnosis. In this study we used the TBD-Serochip to identify peptides that can potentially serve as diagnostic targets for *B. miyamotoi* and differentiate between patients with Lyme disease and BMD.

The most promising candidate diagnostic peptide targets were found on flagella and Vmps, both well-known antigens within relapsing fever *Borreliae*. Flagellar proteins are among the most immunogenic components in spirochetes, highlighted by their inclusion as diagnostic targets in western blot assays for Lyme disease. In previous work, we identified an immunoreactive 13 aa peptide within *B. burgdorferi* FlaB with high diagnostic utility. We subsequently found that this peptide, along with the C6 fragment, were the most frequently reactive *B. burgdorferi* linear peptides in patients with Lyme disease. In addition, we observed that both peptides are often reactive in patients where reactivity to other linear peptides was not detectable. The diagnostic utility of this FlaB fragment extends to *B. miyamotoi,* as the corresponding 14 aa peptide in *B. miyamotoi* was reactive of all samples tested. Although its utility for differential diagnosis is partially diminished by the inability to distinguish between Lyme disease and BMD sera, we found this peptide to be potentially highly useful for diagnosis of *Borrelia* infections, both in early disease (with IgM) or later disease and convalescence (with IgG).

The majority of immunoreactive epitopes identified in our study were located on Vmps (32, 50, 51). Vmps have been shown to be key antigens for neutralization in *Borrelia hermsii* (50). Recent studies in mice have shown the potential of Vsps as a possible target for diagnostic serologic assays for BMD (32). In our study, the majority of reactive peptides were mapped to the variable portions of Vsps and Vlps, but we also identified reactive regions within the conserved protein fragments. Because of higher degree of aa conservation, these reactive peptides could have greater utility for BMD diagnosis.

Our work supports findings from previous studies that reported the C6 ELISA cannot effectively discriminate between antibodies to *B. burgdorferi* and *B. miyamotoi* (20, 26). We also demonstrate that the corresponding C6-like peptide in *B. miyamotoi* is the likely cause of crossreactive signals. Our findings also raise concerns about the specificity of GlpQ as a diagnostic antigen. Three sera that were positive on the GlpQ ELISA did not have reactivity with *B. miyamotoi* peptides in the TBD Serochip assay. We cannot explain the positive commercial ELISA results in the two samples that were negative on the TBD-Serochip and by LIPS. One ELISA-positive sample had a low positive result by LIPS but not the TBD-Serochip. We consider it very unlikely that this patient would have antibodies to *B. miyamotoi* GlpQ but not to any other immunodominant antigens. We conclude that the positive result may have been due to cross-reacting antibodies. Although GlpQ is not present in *B. burgdorferi*, this enzyme is present in a wide range of bacteria, and in some cases, shares substantial aa sequence similarity. For example, the *Escherichia coli* GlpQ shares 49% aa identity with *B. miyamotoi* GlpQ, and in one 39 aa stretch, 34 aa are identical in the two proteins. Thus, it is plausible that in some instances, antibodies to GlpQ from other bacteria may react with *B. miyamotoi* GlpQ ELISA. We suspect that GlpQ specificity may need further examination.

A limitation of our study is that we analyzed a limited number of specimens with confirmed BMD. This was due to the fact that BMD diagnosis is rare and results in a paucity of well-characterized BMD specimens. Also the timing of the convalescent serum samples post treatment might have impacted our findings. Nevertheless, we observed a similar pattern of reactivity from all samples and anticipate that we identified the major reactive *B. miyamotoi* peptides. Although at present the TBD-Serochip is not yet employed for patient serodiagnosis, these peptides can be ported to other serologic platforms that are typically used in clinical laboratories. We utilized a similar approach for the development of a diagnostic assay for Zika virus (52). We anticipate that the panel of peptides we identified in this work can build the foundation of future studies examining the utility of these epitopes for the specific diagnosis of BMD.

## Acknowledgments

We thank Simon H. Williams for his assistance with the manuscript. We thank Sam Telford for providing samples.

This study was funded with grants from the Steven & Alexandra Cohen Foundation (CF CU18-2692 and SACF CU15-4008). This research was supported in part by the Intramural Research Program of the National Institute of Allergy and Infectious Disease, National Institutes of Health.

## Disclaimer

The content of this publication does not necessarily reflect the views of or policies of the Department of Health and Human Services, nor does mention of trade names, commercial products, or organizations imply endorsement by the U.S. Government.

